# YAP-dependent autophagy is controlled by AMPK, SIRT1 and flow intensity in kidney epithelial cells

**DOI:** 10.1101/2023.01.09.523237

**Authors:** Aurore Claude-Taupin, Federica Roccio, Meriem Garfa-Traoré, Alice Regnier, Martine Burtin, Etienne Morel, Fabiola Terzi, Patrice Codogno, Nicolas Dupont

## Abstract

Shear stress generated by the urinary fluid flow is an important regulator of renal function. Its dysregulation is observed in various chronic and acute kidney diseases. Previously, we demonstrated that primary cilium-dependent autophagy allows kidney epithelial cells to adapt their metabolism in response to fluid flow. Here, we show that nuclear YAP/TAZ negatively regulates autophagy machinery in kidney epithelial cells subjected to fluid flow. This crosstalk is supported by a primary cilium-dependent activation of AMPK and SIRT1, independently of the Hippo pathway. We confirmed the relevance of the YAP/TAZ-autophagy molecular dialog *in vivo* using a zebrafish model of kidney development and a unilateral ureteral obstruction mouse model. In addition, an *in vitro* assay simulating the pathological flow observed at early stages of chronic kidney disease (CKD) activated YAP, leading to a primary cilium-dependent inhibition of autophagy. Our findings demonstrate the importance of YAP/TAZ and autophagy in the translation of fluid flow into cellular and physiological responses. Dysregulation of this pathway is associated with the early onset of CKD.

## Introduction

The development and function of organs is tightly controlled by mechanical forces ^1, 2^ In the kidney, mechanical forces, such as shear stress induced by urinary fluid flow, are the principal regulators of proximal reabsorption. Indeed, forces regulate active transport mechanisms in the proximal tubule to drive the reabsorption of up to 70% of Na^+^, K^+^, H^+^, NH_4_^+^, Cl^−^, HCO_3_^−^, Ca^2+^, inorganic phosphate, low-molecular-weight proteins, glucose and water from the glomerular ultrafiltrate into the blood ^3^. Altered tissue mechanics are now recognized to have an active role in driving human diseases ^4–7^. However, how mechanical alterations participate in the early stages of renal diseases, such as chronic kidney disease (CKD), is still unclear.

In kidney proximal tubules, the primary cilium, a microtubule-based organelle present at the apical surface of epithelial cells, acts as a flow sensor to integrate variations in flow rates of the glomerular ultrafiltrate. We and others have demonstrated an interplay between macroautophagy (hereafter referred to as autophagy) and primary cilia ^8–10^. Recently, we showed that primary cilium-dependent autophagy is activated through an LKBI–AMPK signaling pathway to regulate the metabolism and cell volume of kidney epithelial cells ^11, 12^. Autophagy is a catabolic process involved in the maintenance of cellular homeostasis. This self-eating mechanism is initiated by the formation of a double-membrane delimited vacuole, the autophagosome, which ultimately fuses with lysosomes, in order to regulate the turnover of cytoplasmic material, including proteins, lipids and organelles ^13, 14^. Autophagy is tightly coordinated by AuTophaGy-related (ATG) proteins, which can be regulated by transcriptional and post-transcriptional mechanisms^15^.

Yes-associated protein 1 (YAP1; hereafter referred to as YAP) and WW-domain-containing transcription regulator 1 (WWTR1; hereafter referred to as TAZ) are transcriptional coactivators that respond to multiple microenvironmental inputs, including shear stress generated by blood flow in endothelial cells ^16, 17^. YAP/TAZ shuttle between the cytosol and the nucleus where they interact with transcription factors, including the TEA domain family members (TEAD) transcription factors, to induce the expression of key target genes involved in cell proliferation, survival, differentiation or organ size regulation ^18^. The co-transcriptional activity of YAP/TAZ is regulated by the Hippo pathway (consisting of the MST1/2 and LATS1/2 kinases) in response to various signals, such as adhesion proteins or growth factors. Hippo-independent inputs, corresponding to mechanical stimuli such as cell density, stretching or fluid flow also regulate YAP/TAZ activity ^19^. Recently, accumulating evidence has uncovered an interplay between autophagy and the YAP/TAZ pathway ^20–25^. In addition, these pathways have been associated with CKD, such as in diabetic kidney diseases ^25–27^. However, to our knowledge, the crosstalk between autophagy and YAP/TAZ pathway has never been reported in the context of kidney physiology and pathophysiology.

Here, we investigated the crosstalk between these two pathways in the context of shear stress in kidney epithelial cells. We show that YAP/TAZ inactivation by physiological fluid flow is mandatory for the activation of autophagy in proximal tubule kidney epithelial cells. We demonstrate that a primary cilium-dependent activation of AMPK promotes YAP nuclear exit, relying on SIRT1, to inhibit the expression of autophagy repressors, including Rubicon, a suppressor of autophagosome maturation ^28, 29^. Moreover, AMPK-dependent phosphorylation of YAP contributes to its retention in the cytoplasm. Lastly, we show the relevance of this interplay during the early stages of CKD, revealing a potential role of the YAP-autophagy axis dysregulation in renal pathology.

## Results

### YAP/TAZ activity is inhibited by physiological fluid flow

We previously reported that physiological fluid flow stimulates autophagy to direct a metabolic adaptation of proximal tubule kidney epithelial cells (KECs) ^11, 12^. As YAP/TAZ activity can be regulated by different patterns of blood flow ^16^, we investigated whether a physiologically relevant constant laminar flow (1 dyn.cm^-2^) could impact YAP/TAZ activity in KECs. We observed a re-localization of YAP and TAZ into the cytoplasm of KECs submitted to physiological shear stress (Fig. 1a-d). YAP/TAZ are co-transcriptional regulators modulating the activity of TEAD transcription factors. Therefore, we used the 8xGTIIC luciferase reporter, previously described as a YAP/TAZ activity reporter ^30^ to study YAP/TAZ activity upon shear stress. We observed a reduced transactivation activity of YAP/TAZ in cells subjected to fluid flow, as shown by the reduced 8xGTIIC luciferase reporter gene activity (Fig. 1e) and downregulated of the expression of YAP/TAZ target genes (Fig. 1f,g).

**Fig.1:**
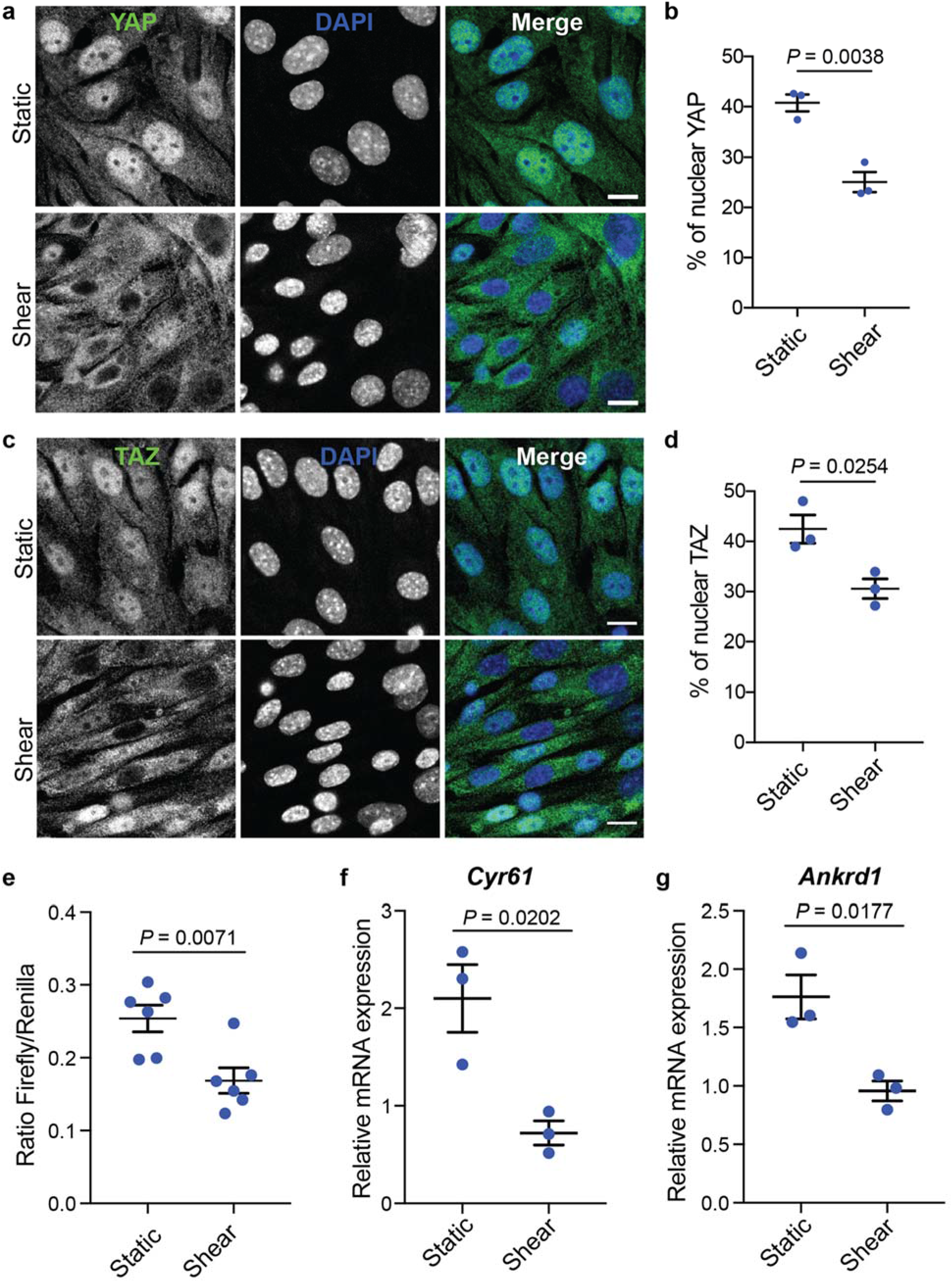
Physiological shear stress inhibits YAP/TAZ. **(a,b)** Representative images **(a)** and quantification **(b)** of YAP localization in the nucleus, labeled with DAPI, in KECs subjected to flow (shear) or not (static) during 24h. Data show the mean ± s.e.m.; n = 3 independent experiments, *t*-test. Scale bars, 10 μm. **(c,d)** Representative images **(c)** and quantification **(d)** of TAZ nuclear localization in KECs subjected to flow (shear) or not (static) during 24h. Data show the mean ± s.e.m.; n = 3 independent experiments, *t*-test. Scale bars, 10 μm. **(e)** Luciferase assay for YAP/TAZ activity in KECs subjected to shear stress or not (static) during 24h. Data show the mean ± s.e.m.; n = 6 from 3 independent experiments, *t*-test. **(f,g)** Expression of *Cyr61* **(f)** and *Ankrd1* **(g)** in KECs subjected to flow (shear) or not (static) during 24h. mRNA levels were quantified by real-time RT-qPCR, normalized to β-actin and are presented as fold increases. Data show the mean ± s.e.m.; n = 3 independent experiments, *t*-test.

As previously described ^24^, YAP/TAZ were mainly localized in the cytoplasm in cells cultivated at high density under static conditions (Extended Data Fig. 1a-d). However, the decrease in YAP/TAZ activity was more important when KECs were subjected to fluid flow than under static conditions. Of note, shear stress does not induce cell proliferation but a reduction in cell size and an increase in cell density ^11, 31^. This suggests that an active mechanism inhibiting YAP/TAZ is dependent on the mechanical forces exerted by unilateral fluid flow. Indeed, the expression of additional YAP/TAZ target genes (*Ptplad2* and *Cptp*), previously reported as being inhibited by fluid flow ^32^, was also downregulated in KECs subjected to shear stress (Extended Data Fig. 1e, f). Thus, under physiological fluid flow conditions, YAP/TAZ are retained in the cytoplasm and are transcriptionally inactive in kidney epithelial cells.

### Autophagy is regulated by YAP/TAZ upon fluid flow

To determine the role of YAP/TAZ activity in fluid flow-induced autophagy, we investigated autophagy in KECs after knocking down the expression of either YAP or TAZ. Of note, the double YAP/TAZ knockdown affected cell viability and proliferation. Physiological fluid flow stimulates autophagy in KECs ^11, 12^, as shown by the increase of endogenous LC3 puncta in cells upon shear stress (Fig. 2a-d). The knockdown of YAP (Fig.2a, b) or TAZ (Fig.2c, d) increased the accumulation of LC3 positive structures in the presence of the lysosomotropic agent chloroquine (CQ), indicating that autophagy flux is amplified in KECs under flow. Accordingly, the accumulation of LC3-II, the lipidated form of LC3 that is associated to autophagosome membranes, was further increasing in cells treated with CQ after YAP or TAZ inactivation, confirming an increase of autophagy flux under these conditions (Extended Data Fig. 2a-d). Thus, even at high density, a condition known to block YAP-dependent autophagy in static conditions (Fig. 2a-d and ^24, 25^), a stimulation of autophagy was observed in KECs submitted to fluid flow. These data suggest that a specific interplay between YAP/TAZ and autophagy occurs when kidney epithelial cells are under physiological shear stress.

**Fig.2:**
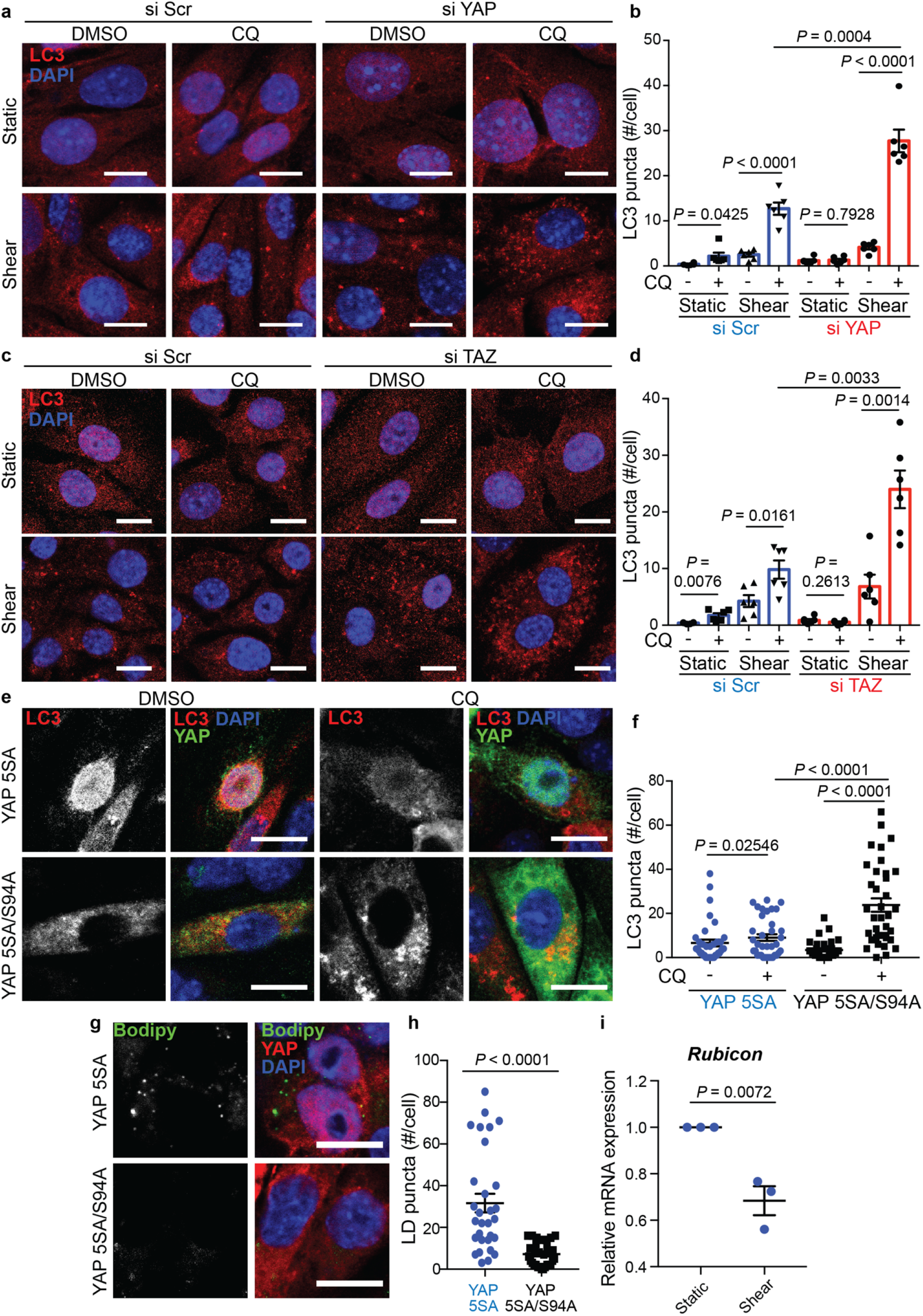
YAP/TAZ regulate autophagy during shear stress. **(a,b)** Representative images **(a)** and quantification **(b)** of LC3 puncta in KECs after transfection with a control siRNA (si Scr) or siRNA against *YAP*, then subjected to flow (shear) or not (static) during 24h in the presence or absence of chloroquine (CQ). Data show the mean ± s.e.m.; n = 6 from 3 independent experiments, *t*-test. Scale bars, 10 μm. #: number. **(c,d)** Representative images **(c)** and quantification **(d)** of LC3 puncta in after transfection with a control siRNA (si Scr) or siRNA against *TAZ*, then subjected to flow (shear) or not (static) during 24h in the presence or absence of chloroquine (CQ). Data show the mean ± s.e.m.; n = 6 from 3 independent experiments, *t*-test. Scale bars, 10 μm. **(e,f)** Representative images **(e)** and quantification **(f)** of LC3 puncta in KECs transfected with the constitutively active form of YAP (YAP 5SA) or inactive form (YAP 5SA/S94A), subjected to shear stress during 24h in the presence or absence of chloroquine (CQ). Data show the mean ± s.e.m.; n = 36 individual data points from 3 independent experiments, *t*-test. Scale bars, 10 μm. **(g,h)** Representative images **(g)** and quantification **(h)** of LDs (Bodipy) in KECs transfected with the constitutively active form of YAP (YAP 5SA) or inactive form (YAP 5SA/S94A), subjected to shear stress during 24h. Data show the mean ± s.e.m.; n = 30 individual data points from 3 independent experiments, *t*-test. Scale bars, 10 μm. **(i)** Expression of *Rubicon* in KECs subjected to flow (shear) or not (static) during 24h. mRNA levels were quantified by real-time RT-qPCR, normalized to β-actin and are presented as fold increases. Data show the mean ± s.e.m.; n = 3 independent experiments, *t*-test.

To confirm the role of YAP/TAZ in the regulation of fluid-flow-dependent autophagy induction, we overexpressed a constitutively active form of YAP (5SA) to counteract YAP inactivation observed in cells submitted to shear stress (see Fig.1). In these conditions, the accumulation of LC3 positive structures was significantly reduced in the presence of CQ, compared to cells overexpressing a constitutively inactive form of YAP (5SA/S94A) (Fig.2 e,f). We obtained similar results when overexpressing a constitutively active form of TAZ (Extended Data Fig. 2e), indicating that YAP/TAZ inhibition by fluid flow is necessary to allow autophagy induction in kidney epithelial cells.

We previously reported that fluid flow induces lipophagy ^12^, a form of selective autophagy that degrades lipid droplets (LDs) ^33^. Therefore, we investigated the role of YAP/TAZ activity on the degradation of LDs in KECs submitted to physiological fluid flow. The expression of YAP 5SA led to the accumulation of LDs upon shear (Fig. 2g, h), supporting the fact that YAP inhibition is required to induce the degradation of LDs in kidney epithelial cells under shear stress.

To better understand how YAP/TAZ regulates autophagy, we studied the expression of *Bcl2* and *Rubicon* ^28, 29, 34^, as these autophagy inhibitors contain TEAD-binding domains in their promotors. We found that the expression of *Rubicon*, but not *Bcl2*, was significantly reduced in cells submitted to fluid flow (Fig. 2i and Extended Data Fig. 2f). In addition, the expression of *Cptp*, another autophagy inhibitor ^35^ but also a YAP/TAZ target gene inhibited by fluid flow ^32^, was reduced in kidney epithelial cells during shear stress (Extended Data Fig.1f). These findings strongly suggest that several autophagy inhibitors are transcriptionally controlled by YAP/TAZ. Hence, shear stress downregulates the expression of YAP/TAZ target genes, including several autophagy inhibitors and allows for autophagy activation.

### The primary cilium is involved in the control of YAP/TAZ activity upon shear stress

We previously reported that kidney epithelial cells need a functional primary cilium (PC) to sense fluid flow and efficiently activate autophagy^11, 12^. To determine the role of PC in regulating the activity of YAP/TAZ upon shear stress, we downregulated the expression of KIF3A, a protein involved in PC assembly^36, 37^ (Extended Data Fig. 3a-c). Downregulation of KIF3A significantly decreased the capacity of YAP/TAZ proteins to exit the nucleus upon shear stress, as observed by immunofluorescence experiments (Fig. 3a-d). Accordingly, we observed an increased transactivation activity of YAP/TAZ in cells depleted for KIF3A, compared to control during shear stress (Fig. 3e). Thus, a functional PC is necessary to efficiently inactivate and retain YAP/TAZ in the cytoplasm.

**Fig.3:**
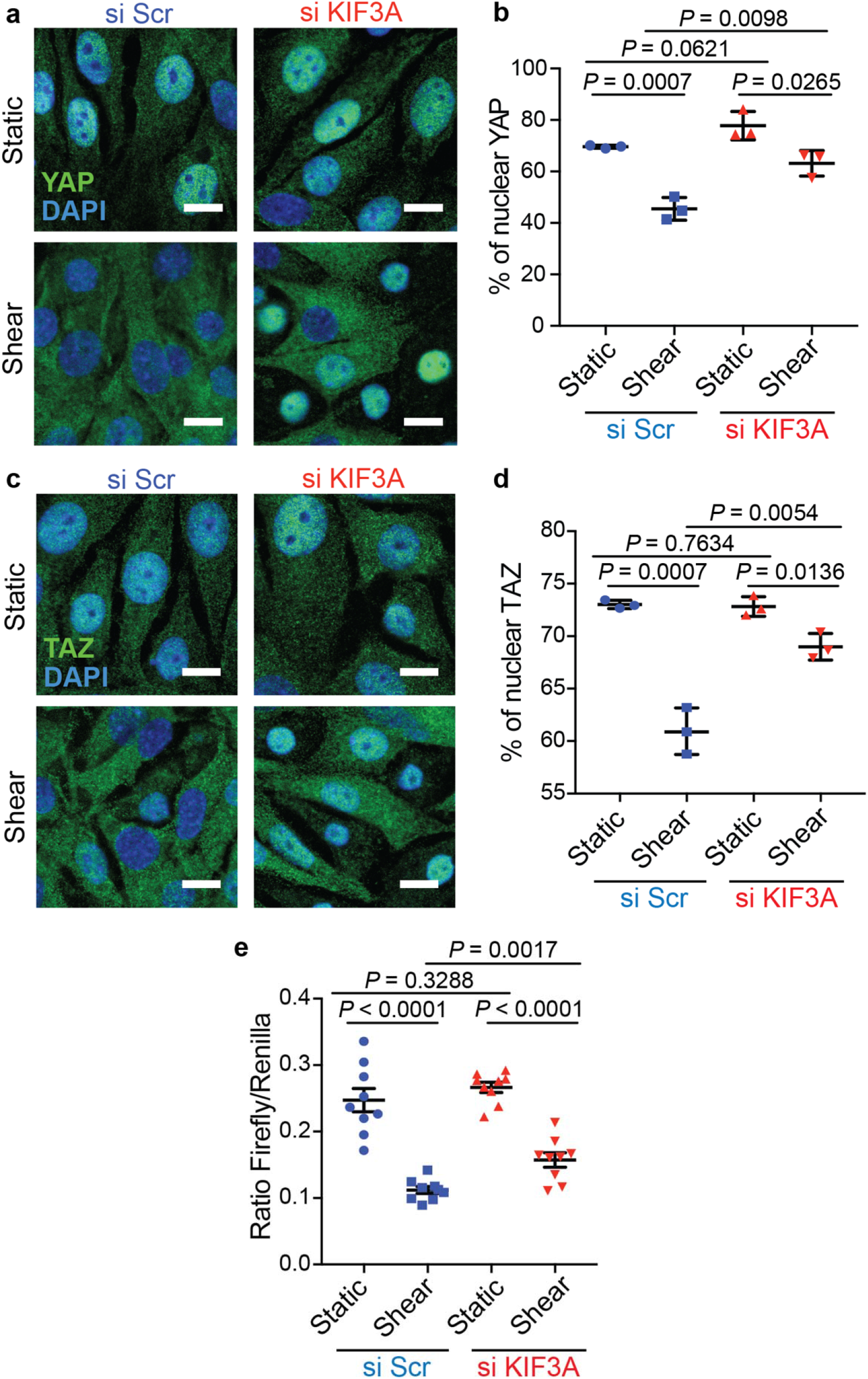
YAP/TAZ inactivation by shear stress requires a functional primary cilium. **(a,b)** Representative images **(a)** and quantification **(b)** of YAP nuclear localization in KECs after transfection with a control siRNA (si Scr) or siRNA against *KIF3A*, subjected to flow (shear) or not (static) during 24h. Data show the mean ± s.e.m.; n = 3 independent experiments, *t*-test. Scale bars, 10 μm. **(c,d)** Representative images **(c)** and quantification **(d)** of TAZ nuclear localization in KECs after transfection with a control siRNA (si Scr) or siRNA against *KIF3A*, subjected to flow (shear) or not (static) during 24h. Data show the mean ± s.e.m.; n = 3 independent experiments, *t*-test. Scale bars, 10 μm. **(e)** Luciferase assay for YAP/TAZ activity in KECs subjected to shear stress or not (static) during 24h, after a knockdown of KIF3A. Data show the mean ± s.e.m.; n = 9 from 3 independent experiments, *t*-test.

### AMPK-dependent regulation of YAP/TAZ by shear stress

The Hippo/LATS kinases can control YAP/TAZ activity by phosphorylation to induce their cytosolic sequestration or degradation ^38^. As we observed cytosolic sequestration of YAP/TAZ upon shear stress, we analyzed the Hippo-dependent phosphorylation of YAP (S127) and TAZ (S89) ^39^ when KECs were subjected to fluid flow (Extended Data Fig. 4a-e). Even though we observed a significant increase of TAZ levels upon shear stress, the ratios of TAZ S89/total TAZ or YAP S127/total YAP remained the same. Accordingly, the levels of active LATS1 (P-LATS1 (T1089)/total LATS1) did not significantly increase during shear stress (Extended Data Fig. 4f,g). Thus, YAP/TAZ inhibition by fluid flow appears to be Hippo-independent.

Upon shear stress, the PC activates autophagy through an LKB1-AMPK signaling pathway ^11, 12^. As YAP can be a substrate of AMPK upon glucose starvation ^40, 41^, we hypothesized that AMPK could phosphorylate YAP upon shear stress. Interestingly, we observed a significant increase in YAP phosphorylation at S61 in KECs subjected to shear stress (Fig. 4a,b). Moreover, YAP phosphorylation at S61 is sufficient to control autophagy induction upon shear stress, as the transfection of YAP S61A inhibited the accumulation of LC3 puncta in the presence of CQ (Fig. 4c,d).

**Fig.4:**
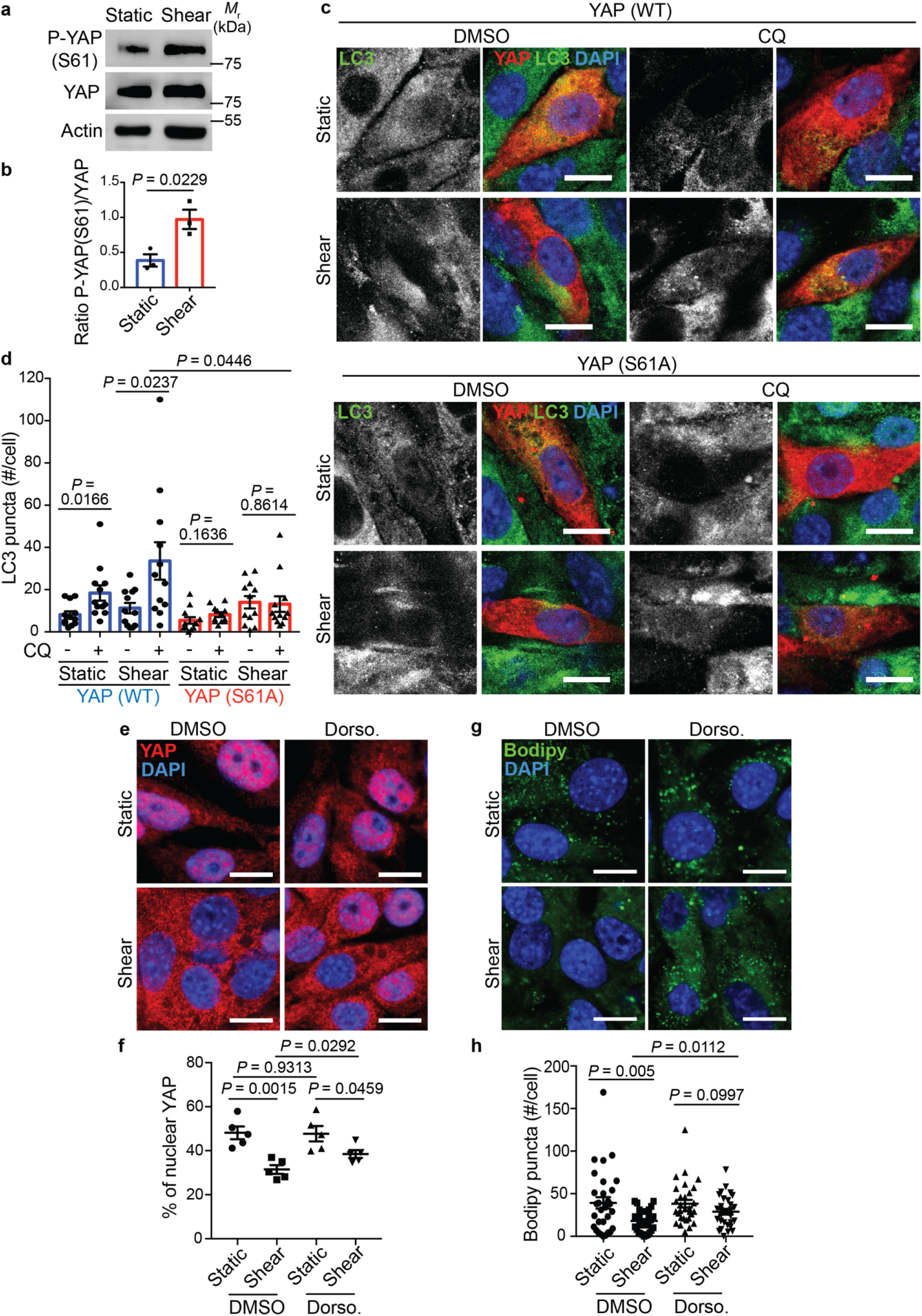
Autophagy is regulated by the AMPK-dependent phosphorylation of YAP at S61 upon fluid flow. **(a)** Representative images of P-YAP (S61), YAP and Actin proteins levels in KECs subjected to flow (shear) or not (static) during 24h, by western blot analysis. **(b)** The ratio of P-YAP (S61) to YAP was determined by densitometry. Data show the mean ± s.e.m.; n = 3 independent experiments, *t*-test. **(c,d)** Representative images **(c)** and quantification **(d)** of LC3 puncta in KECs transfected with YAP WT or YAP S61A mutant, subjected to shear stress during 24h in the presence or absence of chloroquine (CQ). Data show the mean ± s.e.m.; n = 12 individual data points from 3 independent experiments, *t*-test. Scale bars, 10 μm. **(e,f)** Representative images **(e)** and quantification **(f)** of YAP nuclear localization in KECs treated or not with dorsomorphin (Dorso.), subjected to flow (shear) or not (static) during 24h. Data show the mean ± s.e.m.; n = 5 independent experiments, *t*-test. Scale bars, 10 μm. **(g,h)** Representative images **(g)** and quantification **(h)** of lipid droplets (Bodipy) in KECs treated or not with dorsomorphin (Dorso.), subjected to flow (shear) or not (static) during 24h. Data show the mean ± s.e.m.; n = 30 individual data points from 3 independent experiments, *t*-test. Scale bars, 10 μm.

These findings raise the possibility that AMPK-dependent phosphorylation of YAP at S61 could regulate YAP cytoplasmic retention and control autophagy when KECs are cultivated with a physiological stimulus. To confirm this hypothesis, we inhibited the activity of AMPK using dorsomorphin ^42^, which reduced YAP S61 levels (Extended Data Fig.5a). In these conditions, we observed a significant increase of nuclear YAP upon shear stress (Fig. 4e,f) alongside the inhibition of lipophagy (Fig. 4g,h). These data indicate that AMPK-dependent phosphorylation of YAP controls its cytoplasmic retention as well as autophagic processes during shear stress.

### SIRT1 controls YAP exit from the nucleus in response to fluid flow

To better understand the mechanisms responsible for YAP exit from the nucleus upon fluid flow, we studied the potential role of SIRT1 as it has been previously reported as a negative regulator of the co-transcriptional activity of YAP in endothelial cells subjected to laminar flow ^43^. In addition, the deacetylase activity of SIRT1 is dependent on its phosphorylation by AMPK ^44^. Moreover, SIRT1 participates in autophagy activation by controlling the nucleocytoplasmic transport of LC3 upon starvation ^45^. We first analyzed the subcellular localization of SIRT1 and observed an increase of SIRT1 nuclear levels upon shear stress (Fig. 5a,b), suggesting that SIRT1 is activated in these conditions. Indeed, shear stress decreased the levels of the SIRT1 substrate H3K9ac ^46^, while the use of EX527, a SIRT1 inhibitor, preserved H3K9ac levels upon shear (Fig. 5c,d). This suggests that SIRT1 is activated in kidney epithelial cells following shear stress. To investigate its role on YAP translocation to the cytosol upon shear, we inhibited SIRT1 using EX527. SIRT1 inhibition led to the accumulation of nuclear YAP upon shear, as observed by immunofluorescence experiments (Fig. 5e,f). Accordingly, EX527 treatment during shear stress increased the transactivation activity of YAP/TAZ (Extended Data Fig. 5b) and inhibited both autophagy flux (Fig. 5g,h) and LDs degradation (Fig. 5i,j). Previously, we showed that shear stress induced an AMPK-dependent mitochondria biogenesis ^12^. EX527-dependent YAP/TAZ transactivation did not alter the shear stress-dependent expression of *Pgc1a* and *Tfam*, two master genes involved in mitochondria biogenesis (Extended Data Fig. 5c,d). These data suggest that upon shear stress, SIRT1-dependent YAP nuclear exclusion is not involved in the regulation of mitochondria biogenesis but is required for lipo/autophagy induction.

**Fig.5:**
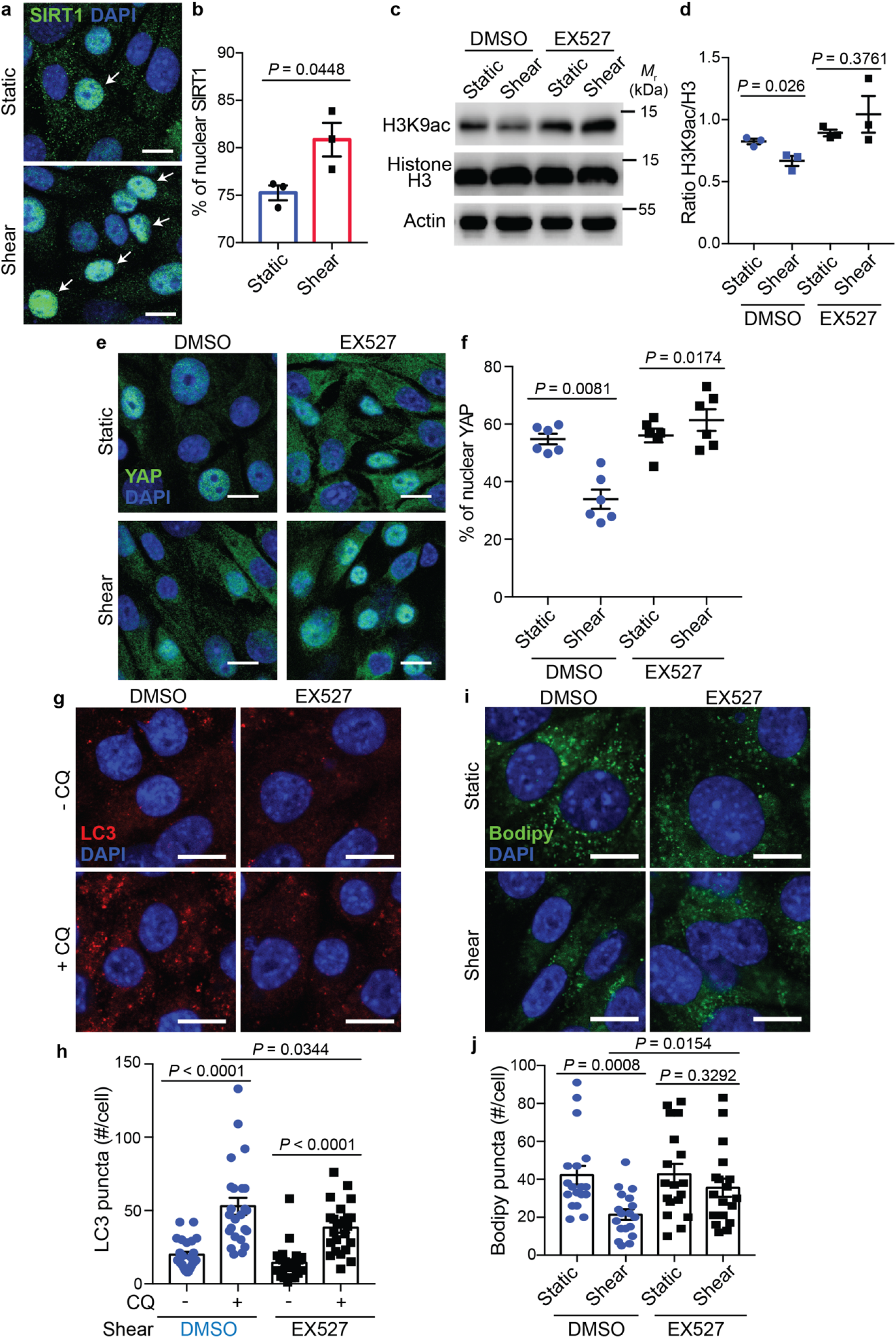
SIRT1 induces YAP nuclear exclusion upon fluid flow. **(a,b)** Representative images **(a)** and quantification **(b)** of SIRT1 nuclear localization in KECs subjected to shear stress during 24h. Data show the mean ± s.e.m.; n = 3 independent experiments, *t*-test. Scale bars, 10 μm. White arrows indicate SIRT1-positive nuclei. **(c)** Representative images of H3K9ac, Histone H3 and Actin proteins levels in KECs subjected to flow (shear) or not (static) during 24h, treated or not with EX527, by western blot analysis. **(d)** The ratio of H3K9ac to Histone H3 was determined by densitometry. Data show the mean ± s.e.m.; n = 3 independent experiments, *t*-test. **(e,f)** Representative images **(e)** and quantification **(f)** of YAP nuclear localization in KECs treated or not with EX527, subjected to flow (shear) or not (static) during 24h. Data show the mean ± s.e.m.; n = 6 individual data points from 3 independent experiments, *t*-test. Scale bars, 10 μm. **(g,h)** Representative images **(g)** and quantification **(h)** of LC3 puncta in KECs subjected to flow (shear) or not (static) during 24h, in the presence or absence of EX527 and chloroquine (CQ). Data show the mean ± s.e.m.; n = 24 individual data points from 3 independent experiments, *t*-test. Scale bars, 10 μm. **(i,j)** Representative images **(i)** and quantification **(j)** of lipid droplets (Bodipy) in KECs treated or not with EX527, subjected to flow (shear) or not (static) during 24h. Data show the mean ± s.e.m.; n = 18 individual data points from 3 independent experiments, *t*-test. Scale bars, 10 μm.

### Effect of the modulation of fluid flow *in vivo* on YAP localization and autophagy

We next aimed to study the relevance of our *in vitro* data using a zebrafish model of kidney development. Indeed, many features of the human nephron are shared with zebrafish pronephros, which begin to filter blood at approximately 48 hours post-fertilization (hpf) ^47^. We analyzed autophagic readouts in the proximal tubule (PT) using wt1b:GFP;LC3:RFP embryos, at 24 hpf when the zebrafish pronephros possess a lumen but no functionality, and at 48 hpf, when the pronephros begins to operate ^47^. We observed that the number of LC3 puncta increased in the PT at 48 hpf compared to 24 hpf (Fig. 6a,b), suggesting that autophagy is induced concomitantly with blood filtration in the kidney pronephros.

**Fig.6:**
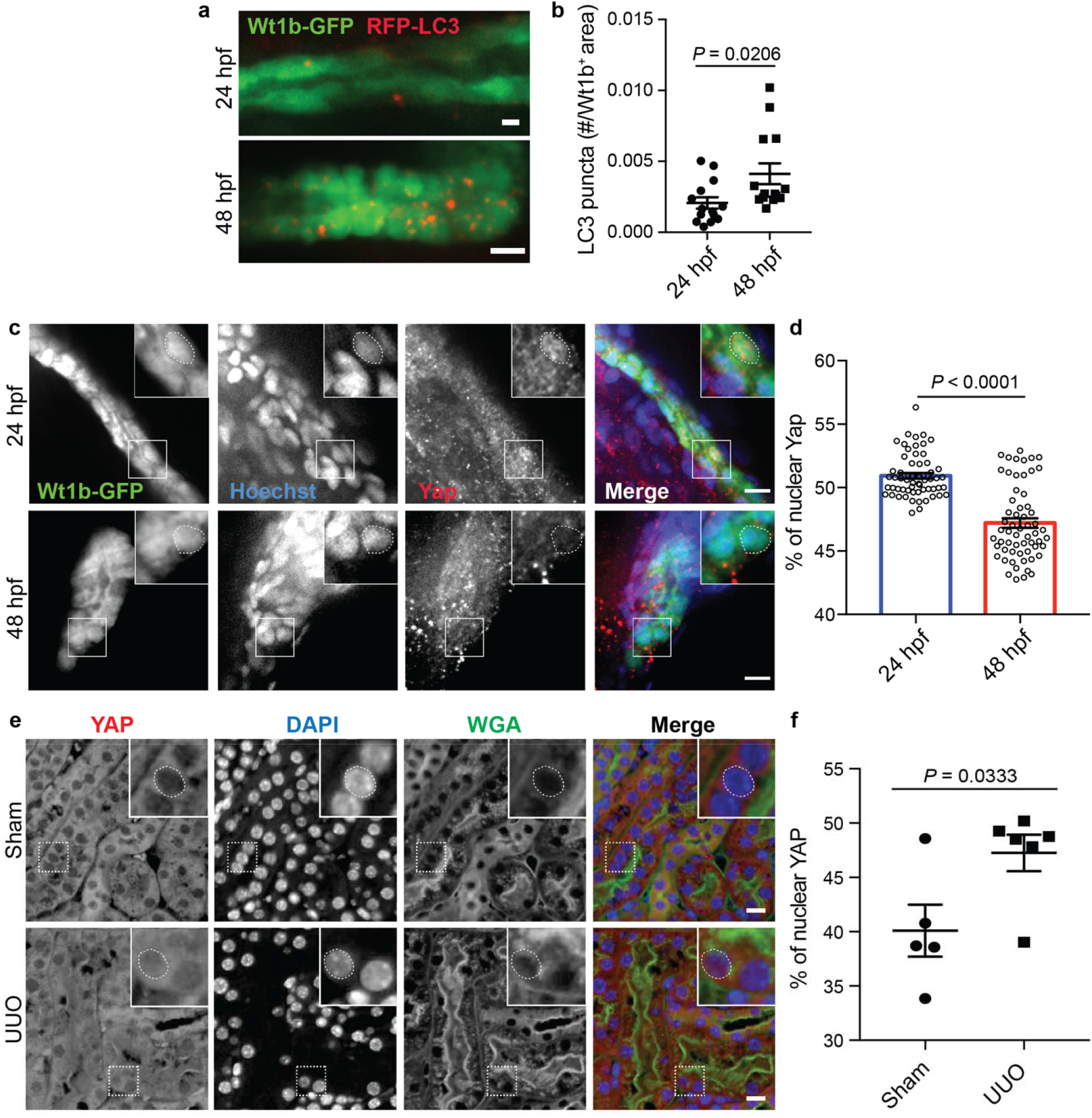
YAP subcellular localization is associated to autophagy activity *in vivo*. **(a,b)** Representative images **(a)** and quantification **(b)** of RFP-LC3 puncta in Wt1b-GFP^+^ pronephros, at 24h post fertilization (hpf) and 48 hpf. Data show the mean ± s.e.m.; n = 18 individual data points from 3 independent experiments, *t*-test. Scale bars, 10 μm. **(c,d)** Representative images **(c)** and quantification **(d)** of YAP nuclear localization in Wt1b-GFP^+^ pronephros at 24 and 48 hpf. Nucleus were labeled with Hoechst. Data show the mean ± s.e.m.; n = 60 individual data points from 3 independent experiments, *t*-test. Scale bars, 10 μm. **(e,f)** Mice subjected to unilateral ureteral obstruction (UUO) or sham operation were euthanized at 24 h after surgery and kidney sections subjected to immunohistochemistry for YAP and renal tubule marker Wheat Germ Agglutinin (WGA). **(e)** Representative images. Scale bars, 10 μm. **(f)** Quantification of YAP nuclear levels in the renal tubules (WGA^+^). Data show the mean ± s.e.m.; n = 5 (sham) and n = 6 (UUO) different mice. *t*-test.

In our *in vitro* model, autophagy induction is dependent on YAP exit from the nucleus. Therefore, we performed immunofluorescence experiments to analyze the subcellular localization of Yap ^48^ at 24- and 48-hpf in the Wt1b^+^ proximal tubules. Using machine learning (see Methods), we delimited the nuclei inside the PT and quantified the percentage of Yap fluorescence in the nuclei and cytoplasm and observed a decrease in the quantity of nuclear Yap at 48-hpf (Fig. 6c,d). Thus, when zebrafish kidney pronephros begin to be functional, Yap translocates to the cytosol and autophagy is induced.

We previously reported that interruption of urinary flow in the kidney of mice by unilateral ureteral obstruction (UUO) resulted in a downregulation of autophagy and accumulation of LDs, at 24h after UUO ^11, 12^. Therefore, we investigated whether YAP subcellular localization in the renal tubule (WGA^+^) could be modified during UUO. We observed a significant increase of nuclear YAP 24h after UUO compared to sham-operated controls (Fig. 6e,f), showing that YAP nuclear translocation is observed at early time points after UUO. This result further confirms an important role of fluid flow in controlling YAP subcellular localization.

Altogether, using two independent *in vivo* models, we confirmed the relevance of the interplay between YAP and autophagy in the functional kidney proximal tubules.

### Pathological fluid flow induces YAP nuclear accumulation and leads to autophagy inhibition

YAP/TAZ dysregulation has recently been linked to CKD ^27, 49^. Upon kidney injury, the loss of functional nephrons triggers compensatory events to maintain kidney function ^50, 51^. Among those events, residual nephrons adapt by increasing their individual nephron filtration levels and urinary fluid flow rates. However, over time, those compensatory events lead to a vicious circle in which the loss of nephrons results in the damage of healthy remaining nephrons. To mimic these pathological conditions, we increased the fluid flow rate from 1 to 4 dyn.cm^2^ (as previously detailed ^52^), using our *in vitro* microfluidic system. In these conditions, a significant increase of nuclear YAP was detected, compared to KECs subjected to a physiological flow condition (1 dyn.cm^-2^) (Fig. 7a,b). Accordingly, the transactivation activity of TEAD was increased in cells subjected to 4 dyn.cm^-2^ flow (Fig. 7c). Of note, at the same time point, TAZ subcellular localization was unchanged (Extended Data Fig. 6a,b), suggesting that an early response of YAP, but not TAZ, is induced by pathological flow. As several studies have demonstrated a protective role of autophagy in renal proximal tubule cells during diabetic nephropathy ^53, 54^, we hypothesized that YAP activation by pathological conditions could be associated with autophagy inhibition. Indeed, autophagy flux was inhibited in cells submitted to high shear stress (Fig. 7d,e). Moreover, it has been shown that under fluid flow, a pool of activated AMPK is present at the primary cilium in kidney epithelial cells ^31^. We observed a decrease of activated AMPK species at the base of primary cilia under high shear stress (4 dyn.cm^-2^) (Fig. 7f,g). Altogether, these data show that there is a causal link between the rate of fluid flow, YAP activity and autophagy dysregulation.

**Fig.7:**
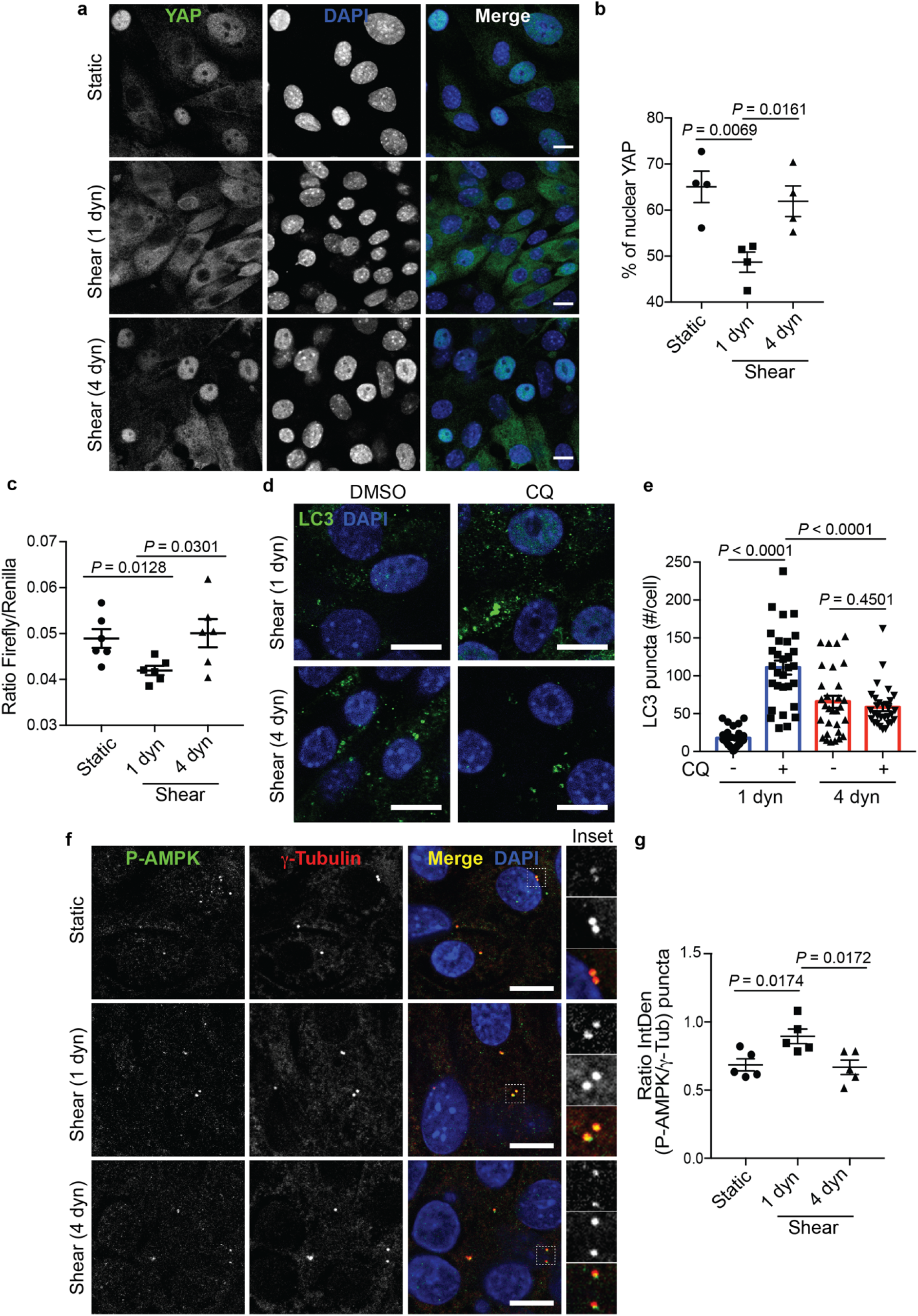
Pathological flow induces YAP nuclear translocation and inhibits autophagy. **(a,b)** Representative images **(a)** and quantification **(b)** of YAP nuclear localization in KECs subjected to physiological flow (shear 1 dyn) or not (static) during 48h. For pathological flow (4 dyn), KECs were subjected to physiological shear stress (1 dyn) during 24h before increasing the flow rate to 4 dyn.cm^-2^ during one more day. Data show the mean ± s.e.m.; n = 4 independent experiments, *t*-test. Scale bars, 10 μm. **(c)** Luciferase assay for YAP/TAZ activity in KECs subjected to shear stress or not (static) during 48h, as described for panel a. 1 dyn = Physiological flow. 4 dyn = pathological flow. Data show the mean ± s.e.m.; n = 6 from 3 independent experiments, *t*-test. **(d,e)** Representative images **(d)** and quantification **(e)** of LC3 puncta in KECs subjected to physiological flow (shear 1 dyn) or pathological flow (shear 4 dyn, as described in panel a), in the presence or absence chloroquine (CQ). Data show the mean ± s.e.m.; n = 30 individual data points from 3 independent experiments, *t*-test. Scale bars, 10 μm. **(f,g)** Representative images **(f)** and quantification **(g)** of colocalization between phosphorylated AMPK (P-AMPK) and centrioles (γ-Tubulin) in KECs subjected to physiological flow (shear 1 dyn), pathological flow (shear 4 dyn, as described in panel a), or not (Static). Data show the mean ± s.e.m. of P-AMPK intensity, compared with γ-tubulin intensity; n = 5 independent experiments, *t*-test. Scale bars, 10 μm.

## Discussion

In this study, we uncovered molecular mechanostranduction mechanisms connecting YAP/TAZ and autophagy in kidney epithelial cells under shear stress. This process requires the activation of AMPK which allows efficient inhibition of YAP/TAZ transcriptional activity, upstream of the expression of autophagy inhibitors CPTP and Rubicon. This transcriptional regulation is achieved by a direct phosphorylation of YAP on S61 by AMPK and by AMPK- and SIRT1-dependent YAP nuclear exclusion (Extended Data Fig.7). While AMPK directly triggers mitochondrial biogenesis ^12^, the metabolic adaptation of proximal tubule cells to shear stress relies on YAP to promote lipophagy, thus supporting energy-consuming cellular processes such as glucose reabsorption and neoglucogenesis ^12^. Altogether, our data and previous results from this laboratory ^11, 12^ and others ^31^, ^55^ demonstrate that the activation of AMPK plays a central role in maintaining the homeostasis of renal tubular cells, which constantly face physiological fluid flow.

Here, we also reported that the PC is important to drive fluid flow-dependent YAP/TAZ nuclear exclusion. AMPK activity at the PC is dependent on the upstream kinase LKB1 ^11^. However, how PC control the activation of LKB1-AMPK axis is still unclear. Based on literature showing the interaction between polycystin-1 (PC1) and LKB1 ^56^, it can be hypothesized that this molecular pathway could be a good candidate to control AMPK activity in KECs subjected to fluid flow.

YAP/TAZ nuclear exclusion is a crucial step to understand the role of the PC-autophagy axis in the control of cell homeostasis. We reported that YAP/TAZ exit from the nucleus requires SIRT1 activity. However, other proteins, such as XPO1 (known as exportin-1), previously shown to regulate YAP nucleo-cytoplasmic shuttling ^57^, could also participate in the control of YAP/TAZ localization. Upon shear stress, the cytoplasmic accumulation of YAP/TAZ decreases the levels of *Rubicon* to control autophagy and metabolism. Indeed, a knockdown of YAP/TAZ led to high autophagic flux levels. These results are in agreement with a recent study which showed that Rubicon-deficient kidney proximal tubular cells exhibit high autophagy flux and accelerated LDs degradation ^58^, suggesting the importance of balancing the activity of YAP/TAZ and thus autophagy, to maintain kidney tubular cells homeostasis.

In this study, we revealed the importance of mechanical stimuli to control YAP/TAZ and autophagy. Previous reports have shown that matrix stiffness ^25^ and contact inhibition ^23, 24^ are important factors regulating the YAP/TAZ-autophagy axis. In these contexts, autophagy is compromised upon YAP/TAZ inhibition (cytoplasm sequestration) and we confirmed this in kidney epithelial cells cultivated at high density under static conditions (See Fig. 2). However, simulating physiological shear stress *in vitro* on cells at high density stimulated autophagy in a cytoplasmic YAP/TAZ-dependent manner. Interestingly, Hippo-dependent cytoplasmic retention of YAP/TAZ induced by high density or matrix stiffness, inhibits autophagy by repressing the expression of proteins engaged in the formation ^23, 24^ and maturation ^25^ of autophagosomes. In this work, we show that under shear stress, a Hippo-independent cytoplasmic retention of YAP/TAZ stimulates autophagy by repressing the expression of autophagy inhibitors (*Rubicon* and *Cptp*). It should be kept in mind that the different cytoplasmic forms of phospho-YAP could recruit different partners ^59, 60^, to control autophagy in a posttranscriptional manner. Thus, according to the mechanical stress sensed by the cell, a specific response will occur to translate these forces into the appropriate metabolic adaptation.

AMPK and autophagy are well known to be dysregulated in renal diseases, including in CKD ^61^. In addition, several GWAS studies of CKD have identified several loci, including a locus containing *PRKAG2*, the gene encoding the AMPK γ2 subunit. Of note, this gene has been shown to be associated with rapid kidney function decline ^62, 63^. Based on this literature, different molecules that directly or indirectly activate AMPK, such as metformin (already widely used to treat type 2 diabetes mellitus), have been identified as efficient inhibitors of the progression of renal diseases, including polycystic kidney disease, renal cancer or acute kidney injury ^61, 64^. Despite these promising results, the use of metformin is not easily translatable to humans, since its use is not devoid of side effects, such as lactic acidosis. Here, we reported that a pathological fluid flow mimicking the shear stress observed in CKD is sufficient to re-localize YAP into the nucleus and block the autophagic pathway. These data are in line with previous studies showing that YAP is hyperactivated in several CKD mouse models and in CKD patients ^65^. More importantly, we have shown that those effects are correlated with a defect of AMPK activation at the base of the primary cilia. Thus, our study has shed light on the role of a specific ciliary pool of AMPK in maintaining the homeostasis of renal tubular cells and opens new avenues to develop therapeutical strategies for CKD patients.

---

**Extended Data Fig.1:**
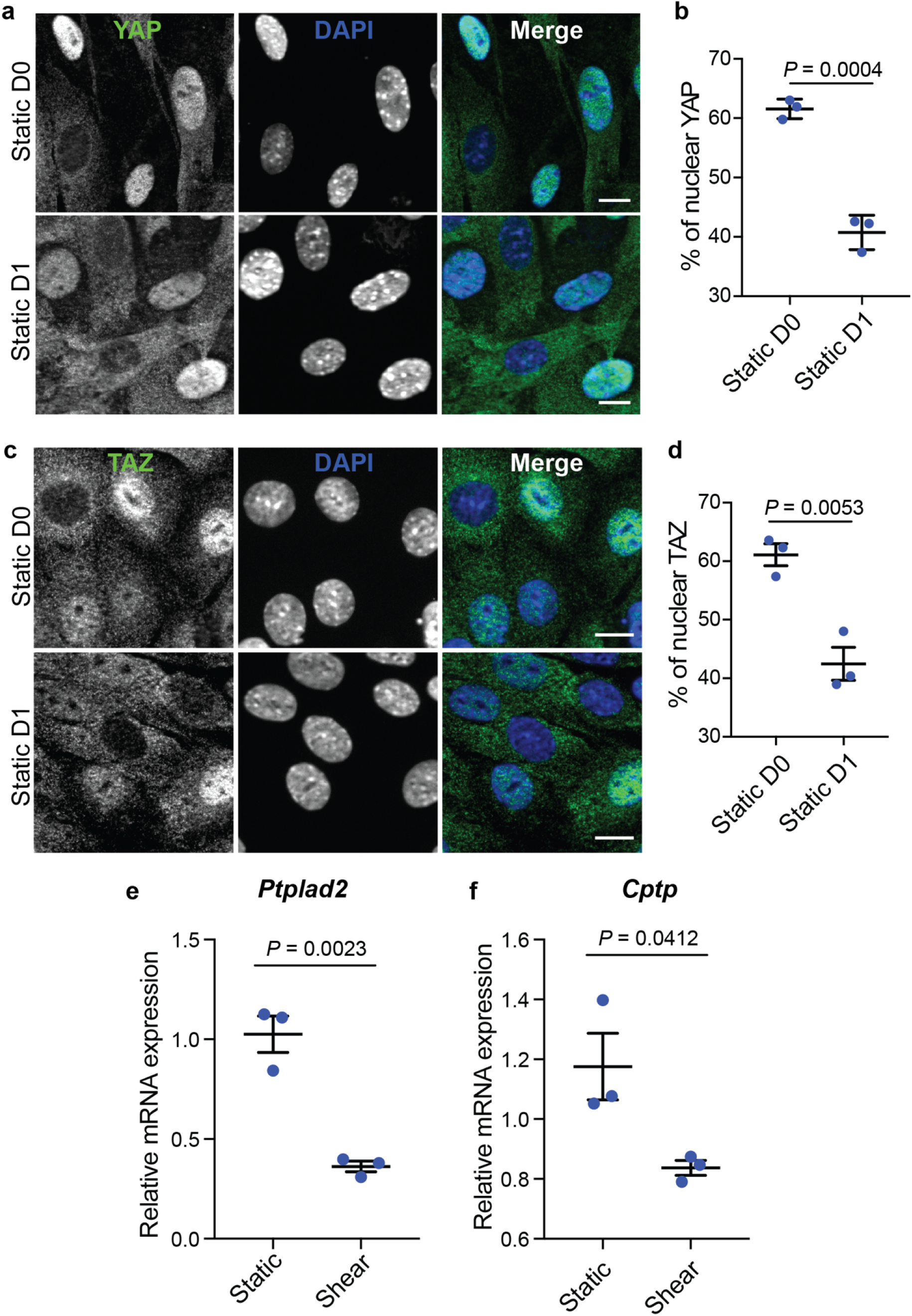
YAP/TAZ inhibition by shear stress leads to the downregulation of flow-specific YAP/TAZ target genes. **(a,b)** Representative images **(a)** and quantification **(b)** of YAP nuclear localization in KECs plated at high density (Static D0) and kept in culture for 24h (Static D1). Data show the mean ± s.e.m.; n = 3 independent experiments, *t*-test. Scale bars, 10 μm. **(c,d)** Representative images **(c)** and quantification **(d)** of TAZ nuclear localization in KECs plated at high density (Static D0) and kept in culture for 24h (Static D1). Data show the mean ± s.e.m.; n = 3 independent experiments, *t*-test. Scale bars, 10 μm. **(e,f)** Expression of *Ptplad2* **(e)** and *Cptp* **(f)** in KECs subjected to flow (shear) or not (static) during 24h. mRNA levels were quantified by real-time RT-qPCR, normalized to β-actin and are presented as fold increases. Data show the mean ± s.e.m.; n = 3 independent experiments, *t*-test.

**Extended Data Fig.2:**
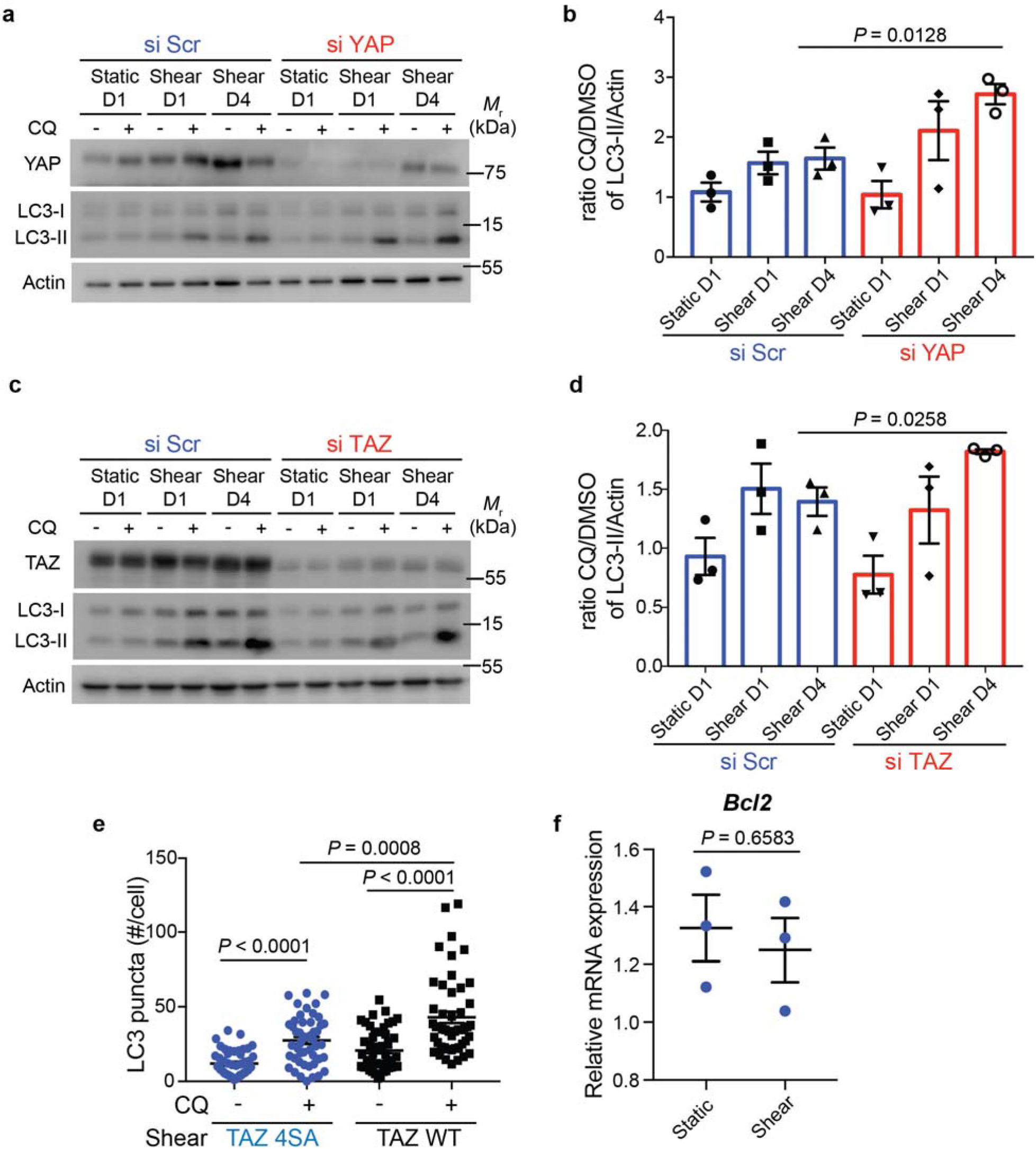
YAP and TAZ control autophagy during shear stress. **(a)** Representative images of YAP, LC3-I, LC3-II and Actin proteins levels in KECs after transfection with a control siRNA (si Scr) or siRNA against *YAP*, subjected to flow (shear, during 1 or 4 days) or not (static), in the presence or absence of chloroquine (CQ), by western blot analysis. **(b)** The ratio of LC3-II to Actin was determined by densitometry, relative to panel a. Data show the mean ± s.e.m.; n = 3 independent experiments, *t*-test. **(c)** Representative images of TAZ, LC3-I, LC3-II and Actin proteins levels in KECs after transfection with a control siRNA (si Scr) or siRNA against *TAZ*, subjected to flow (shear, during 1 or 4 days) or not (static), in the presence or absence of chloroquine (CQ), by western blot analysis. **(d)** The ratio of LC3-II to Actin was determined by densitometry, relative to panel c. Data show the mean ± s.e.m.; n = 3 independent experiments, *t*-test. **(e)** Quantification of LC3 puncta in KECs transfected with TAZ WT or its constitutively active form (TAZ 4SA), subjected to shear stress during 24h in the presence or absence of chloroquine (CQ). Data show the mean ± s.e.m.; n = 48 individual data points from 3 independent experiments, *t*-test. **(f)** Expression of *Bcl2* in KECs subjected to flow (shear) or not (static) during 24h. mRNA levels were quantified by real-time RT-qPCR, normalized to β-actin and are presented as fold increases. Data show the mean ± s.e.m.; n = 3 independent experiments, *t*-test.

**Extended Data Fig.3:**
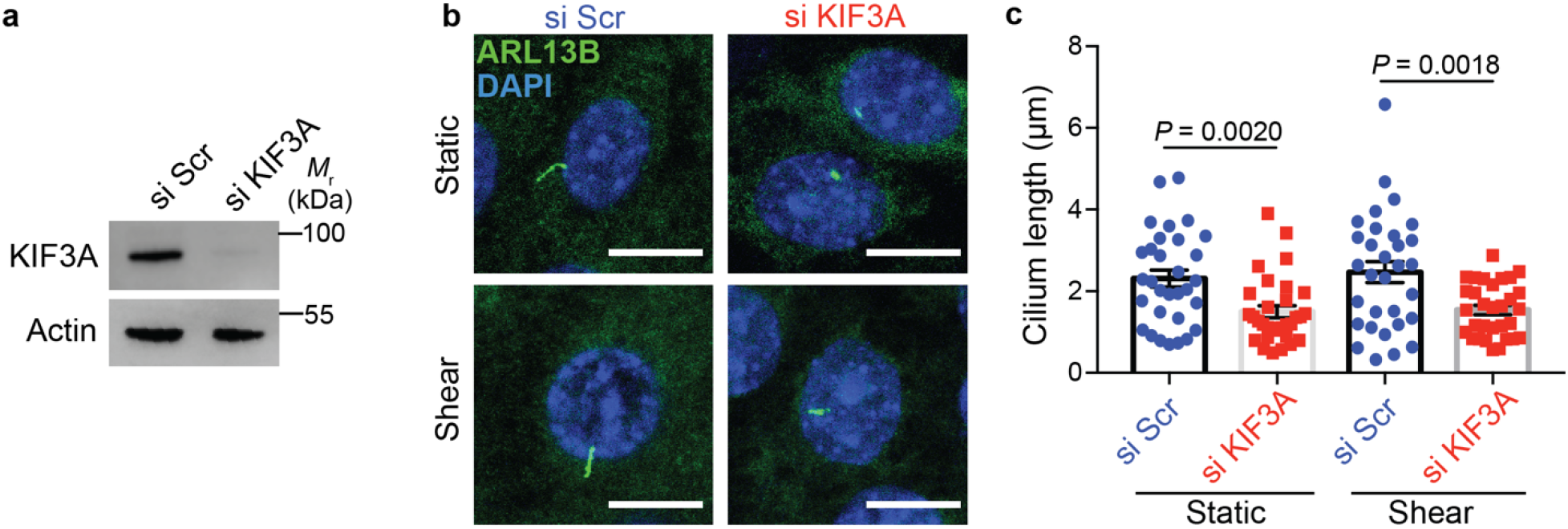
KIF3A downregulation in KECs affects ciliogenesis. **(a)** Confirmation by immunoblotting of KIF3A knockdown in KEC cells. Representative image from n=3 independent experiments shown. **(b,c)** Representative images **(b)** and quantification **(c)** of cilia length (ARL13B^+^) in KECs after transfection with a control siRNA (si Scr) or siRNA against *KIF3A*, subjected to flow (shear) or not (static) during 24h. Data show the mean ± s.e.m.; n = 31 individual data points from 3 independent experiments, *t*-test. Scale bars, 10 μm.

**Extended Data Fig.4:**
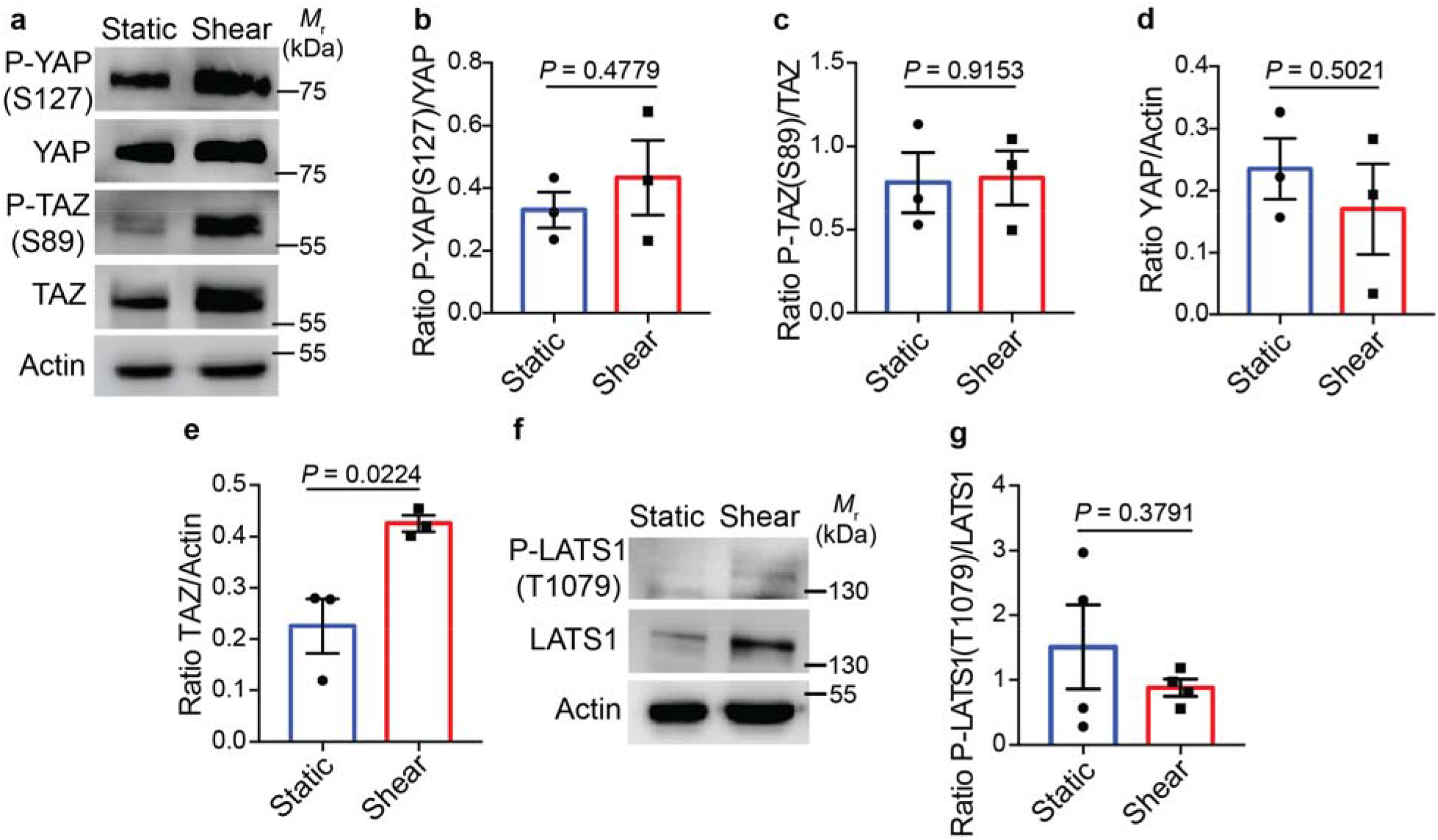
YAP/TAZ inactivation during shear stress is Hippo independent. **(a)** Representative images of P-YAP (S127), YAP, P-TAZ (S89), TAZ and Actin proteins levels in KECs subjected to flow (shear) or not (static) during 24h, by western blot analysis. **(b-e)** The ratio of P-YAP to YAP **(b)** P-TAZ to TAZ **(c)** YAP to Actin **(d)** and TAZ/Actin **(e)** was determined by densitometry, relative to panel a. Data show the mean ± s.e.m.; n = 3 independent experiments, *t*-test. **(f)** Representative images of P-LATS1 (T1079), LATS1 and Actin proteins levels in KECs subjected to flow (shear) or not (static) during 24h, by western blot analysis. **(g)** The ratio of P-LATS1 to LATS1 was determined by densitometry, relative to panel f. Data show the mean ± s.e.m.; n = 3 independent experiments, *t*-test.

**Extended Data Fig.5:**
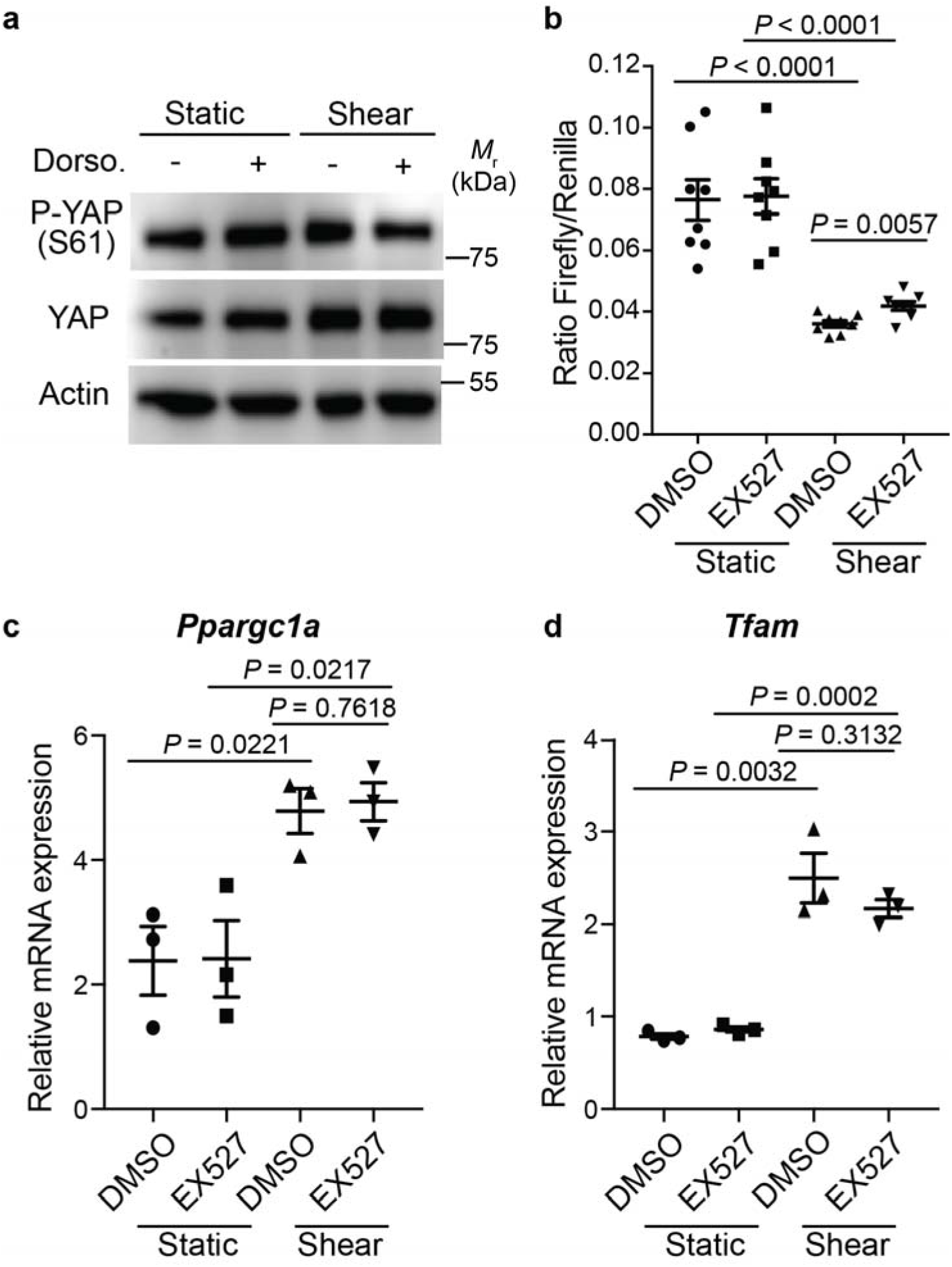
YAP cytoplasmic retention, dependent on AMPK and SIRT1, does not influence mitochondrial biogenesis. **(a)** Representative images of P-YAP (S61), YAP and Actin proteins levels in KECs subjected to flow (shear) or not (static) during 24h, in the presence or absence of dorsomorphin (Dorso.) by western blot analysis. **(b)** Luciferase assay for YAP/TAZ activity in KECs subjected to shear stress or not (static) during 24h, in the presence or absence of EX527. Data show the mean ± s.e.m.; n = 8 from 4 independent experiments, *t*-test. **(c,d)** Expression of *Ppargc1a* **(c)** and *Tfam* **(d)** in KECs subjected to flow (shear) or not (static) during 24h, in the presence or absence of EX527. mRNA levels were quantified by real-time RT-qPCR, normalized to β-actin and are presented as fold increases. Data show the mean ± s.e.m.; n = 3 independent experiments, *t*-test.

**Extended Data Fig.6:**
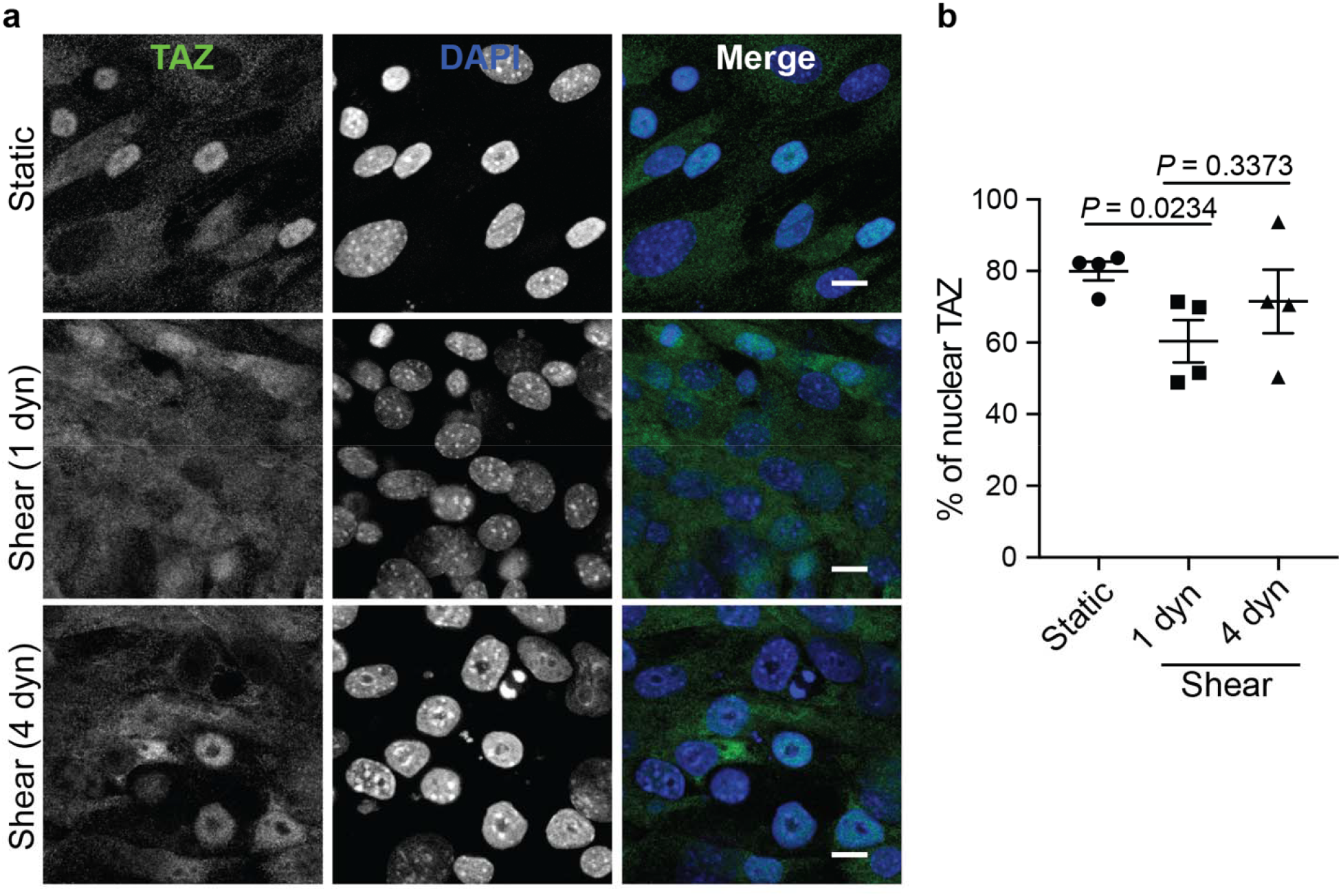
Pathological flow does not induce TAZ nuclear translocation. Representative images **(a)** and quantification **(b)** of TAZ nuclear localization in KECs subjected to physiological flow (shear 1 dyn) or not (static) during 48h. For pathological flow (4 dyn), KECs were subjected to physiological shear stress (1 dyn) during 24h before increasing the flow rate to 4 dyn.cm^-2^ during one more day. Data show the mean ± s.e.m.; n = 4 independent experiments, *t*-test. Scale bars, 10 μm.

**Extended Data Fig.7:**
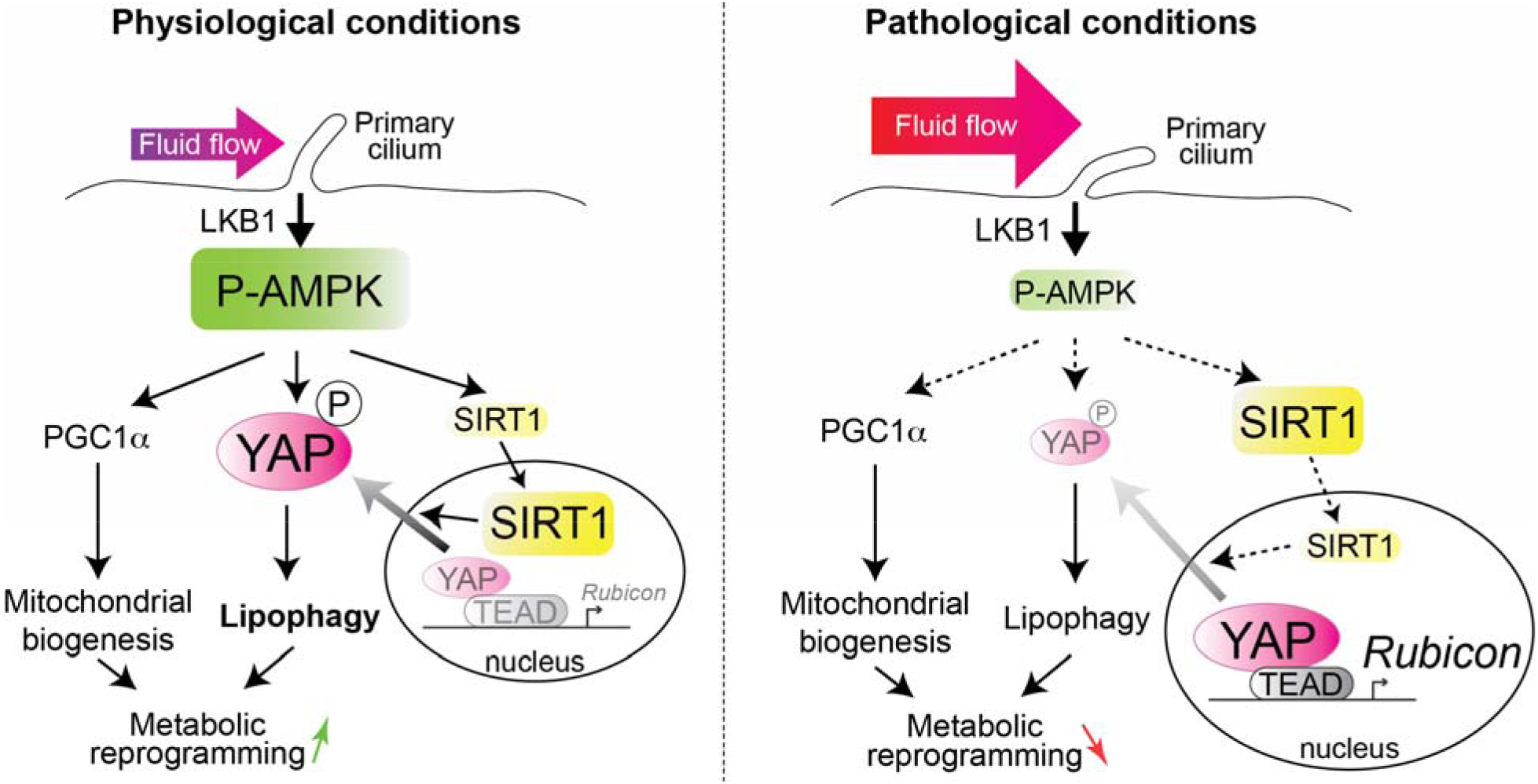
Overall schematic summary. In physiological conditions, the primary cilium-dependent activation of AMPK ensures the stimulation of 3 pathways: (i) SIRT1 activation is necessary to induce YAP exit from the nucleus (ii) AMPK-dependent phosphorylation of YAP on S61 sequestrate YAP in the cytosol, thus inhibiting the transcription of *Rubicon*, an autophagy suppressor. (iii) Increase of mitochondrial biogenesis. This metabolic reprogramming supports energy-consuming cellular processes such as glucose reabsorption and gluconeogenesis. In pathological conditions, the primary cilium-dependent activation of AMPK is impaired, resulting notably in the accumulation of nuclear YAP and inhibition of autophagy.

## Methods

### Cell culture, transfections and plasmids

Wild-type mouse KECs, generated by G.J. Pazour (University of Massachusetts), were kindly provided by A.M. Cuervo (Albert Einstein College). The cells used in this study were free of mycoplasma contamination. No cell lines used in this study were found in the database of commonly misidentified cell lines maintained by ICLAC and NCBI Biosample.

To inhibit the expression of YAP, TAZ and KIF3A, cells were doubly transfected using RNAiMAX (Invitrogen) according to the manufacturer’s instructions with siRNAs targeting mRNAs encoding for YAP, TAZ or KIF3A from Qiagen (listed in Supplementary Table 1). After transfection, cells were seeded onto microslides and subjected to shear stress the following day. Non-targeting siRNA pool (Scramble) was used as a control.

DNA transfections were done with Lipofectamine 2000 (Invitrogen). Plasmids used in this study are listed in Supplementary Table 2. YAP S61A mutant was generated utilizing a QuikChange site-directed mutagenesis kit (Agilent) and confirmed by sequencing (Eurofins Genomics).

### Flow chamber

KECs were subjected to shear stress as previously described ^11^. Briefly, cells were seeded in closed perfusion chambers (Microslide I^0.6^ Luer; channel dimensions 50 × 5 × 0.4 mm coated with iBiTreat, Ibidi) and cultured for at least one day at full confluency to allow polarization and maximum ciliogenesis. Medium was changed twice a day before flow induction. The chamber was then connected to a computer-controlled set-up containing an air-pressure pump and a two-way switching valve (Ibidi pump system #10902) using a perfusion set (Ibidi Blue #10961, length 15 cm, ID 0.8 mm). Ten mL of cell culture medium was pumped unidirectionally between two reservoirs through the flow channel at a rate corresponding to a shear stress of 1 dyn.cm^-2^ (physiological fluid flow). To simulate pathological fluid flow, cells were subjected to 1 dyn.cm^-2^ shear stress for 1 day to allow KECs’ differentiation. Then, the fluid flow was increased to 4 dyn.cm^-2^ for 1 day before analysis, to simulate the flow rate observed soon after subtotal nephrectomy by micropuncture studies ^50, 66^.

### Pharmacological inhibitors

All autophagic measurements and treatments were performed as described previously ^67^. Briefly, where indicated, cells were treated with chloroquine (CQ, 10 μM, Sigma-Aldrich) to inhibit autophagic maturation. To inhibit AMPK and SIRT1, cells were respectively treated with dorsomorphin (Dorso., 10 μM, Sigma-Aldrich) or EX527 (10 μM, Sigma-Aldrich).

### Immunoblotting, antibodies and detection assays

Blots were labelled with antibodies against LC3B (Sigma, #L7543, 1:5000 dilution), YAP (Cell Signaling Technology, #4912, 1:1000), Phospho-YAP_S127 (Cell Signaling Technology, #13008, 1:1000), Phospho-YAP_S61 (Cell Signaling Technology, #75784, 1:1000), TAZ (Sigma, #HPA007415, 1:1000), Phospho-TAZ_S89 (Cell Signaling Technology, #59971, 1:1000), LATS1 (Cell Signaling Technology, #9153, 1:1000), Phospho-LATS1_T1079 (Cell Signaling Technology, #8654, 1:1000), H3K9ac (Sigma, #07-352, 1:1000), Histone H3 (Proteintech, #17168-1-AP, 1:1000) and actin (Millipore, Clone C4 MAB1501, 1:20,000 dilution). Appropriate HRP-labelled anti-rabbit (Millipore, #AP307P, 1:10000) and anti-mouse (Millipore, #AP308P, 1:10000) were then used and revealed with Super Signal West Dura chemiluminescent substrate (Pierce). Images were taken with the ChemiDoc-MP Imaging System and quantified using ImageJ software.

### qRT–PCR

Total RNA was extracted from cells using the Cells-to-Ct Kit (Applied Biosystems), according to the manufacturers’ instructions. Real-time PCR was performed using SYBR Green Master Mix (Applied Biosystems), and products were detected on a qTOWER 2.0 Real-Time PCR System (Analytik Jena). Relative expressions of *Cyr61, Ankrd1, Rubicon, Ptplad2, Cptp, Bcl2* and *β-actin* were calculated using the 2^ΔΔC(t)^ method. *β*-actin was used as normalization control. Conditions for real-time PCR were as follows: initial denaturation for 10 min at 95 °C, followed by amplification cycles of 15 s at 95 °C, and 1 min at 60 °C. The qPCR primers used in this study are listed in Supplementary Table 3.

### Conventional microscopy and antibodies

Immunofluorescence microscopy was carried out using a Zeiss Confocal LSM700 piloted with Zen software. Cells grown on microslides were fixed for 10 min at RT in 4% paraformaldehyde (PFA), permeabilized (5 min, 0.1% Triton X-100), blocked (30 min, 10% FBS) and incubated with the following primary antibodies in blocking buffer for 2 hours at RT: YAP (Cell Signaling Technology, #14074, 1:100), TAZ (Sigma, #HPA007415, 1:100), LC3B (MBL, #PM036 or #M152.3, 1:200 dilution), FLAG-M2 (Sigma, # F1804, 1:200), followed by incubation with fluorophore-conjugated secondary antibodies (Molecular Probes) for 1 hour. For primary cilia imaging, cells were fixed with cold methanol for 10 min at −20°C, blocked (30 min, 10% FBS) and incubated with the following primary antibodies in blocking buffer containing 0.05% Saponin for 2 hours at RT: ARL13B (Proteintech, # 66739, 1:100), γ-tubulin (Sigma, #T5326, 1:200), Phospho-AMPK_T172 (Cell Signaling Technology, #2535, 1:200), followed by incubation with fluorophore-conjugated secondary antibodies (Molecular Probes) for 1 hour.

The cells were also labelled with Bodipy 493/503 (Thermo Fisher Scientific, #D3922, 1:200) to detect the LDs by fluorescence microscopy, as described previously ^68^. Mounting medium containing 4’,6-diamidino-2-phenylindole (DAPI) was used when required. ImageJ software was used to quantify the percentage of nuclear YAP/TAZ (Intensity Ratio Nuclei Cytoplasm Tool, RRID:SCR_018573) and colocalization between Phospho-AMPK and γ-tubulin (Red and Green Puncta colocalization Macro, D. J. Swiwarski modified by R.K. Dagda). Icy software was used to quantify LC3 and Bodipy puncta, and cilia length.

### Luciferase reporter assay

The luciferase reporter construct 8xGTIIC-Luciferase (Firefly) was a gift from Stefano Piccolo (Addgene plasmid #34615) ^30^. KECs were transfected with 8xGTIIC plasmid along with Renilla luciferase (Ratio 10:1). The next day, cells were seeded onto microslides and subjected to shear stress the following day. At the end of shear stress, luciferase activity was measured using Pierce Renilla-Firefly Luciferase Dual Assay Kit (Thermo Scientific, #16185), according to manufacturer’s instructions.

### Animals and protocols

Mice were on a C57BL/6 genetic background (Janvier Laboratories). Animal procedures were approved by the departmental director of the Services Vétérinaires de la Préfecture de Police de Paris and by the ethical committee of the Paris Descartes University. Animals were housed in a specific pathogen-free facility, fed *ad libitum*, and housed at constant ambient temperature and humidity ranges (50–60%) in a 12h light cycle. Nine-week-old female mice were subjected to UUO (n = 6) or sham operation (controls; n = 5) as previously described ^69^ and euthanized 24 h after surgery. During the same period (24 h), mice were deprived of food. At euthanasia, kidneys were removed for immunofluorescence analysis. Briefly, 4-μm sections of paraffin-embedded kidneys were submitted to the appropriate antigen retrieval. Then, sections were incubated with YAP (Cell Signaling Technology, #14074, 1:200) antibody overnight at 4°C, followed with the appropriate secondary antibody and WGA-Rhodamine (Vector Laboratories, RL-1022, 1:200). DAPI was used to stain nuclei. Whole kidney sections were then scanned using a nanozoomer 2.0 HT (Hamamatsu) with a ×40 immersion objective. ImageJ software was used to quantify the percentage of nuclear YAP (Intensity Ratio Nuclei Cytoplasm Tool, RRID:SCR_018573) in the proximal kidney tubules.

The Zebrafish Tg(hsp70l:RFP-Rno.Map1lc3b) line (RFP-LC3) was obtained from Dr. Enrico Moro (University of Padova, Italy) ^70^ and the Tg(wt1b:GFP) line ^71^ was used to label proximal pronephros. Embryos were obtained by natural mating and raised at 28.5°C in petri dishes containing fish water. To induce the expression of RFP-LC3, larvae were collected 6 hpf and heat shocked by placing in 42°C fish water and then incubating for 60 min at 37°C. Fish were then placed back at 28.5°C until live imaging at 24 and 48 hpf. To avoid pigmentation of embryos, 0.003% 1-phenyl-2-thiourea (PTU) was added to the embryo medium 24 hpf. Immunofluorescence experiments were performed as described previously ^72^. Briefly, embryos were fixed overnight at 4°C in 4% PFA, then washed 3 times for 20 min in PBS 1% Triton X-100 (PBS-Triton). Embryos were incubated for 1 hr in blocking buffer (PBS-Triton, FBS 10%) then overnight at 4°C with anti-Yap antibody (Cell Signaling, #4912, 1:200) in blocking buffer. Embryos were then washed 3 times for 20 min in PBS-Triton before incubation with anti-rabbit Alexa Fluor 546 used at 1:500 dilution in blocking buffer containing Hoechst 33342 (Sigma) to label the nuclei. After 3 washes in PBS-Triton, embryos were mounted in 2% low melting agarose. Z-stacks were acquired using a Zeiss Spinning disk. For image analysis, tubule volume and nucleus were first segmented using Ilastik pixel classification (v1.3.3post3) ^73^. Then with a Fiji (v2.3.0/1.53f51) ^74^ macro, we manually corrected the tubule volume and selected several ROIs for mean background measurements, to finally measure the ratio of the mean tubular YAP intensity, with background subtraction, inside and outside the nucleus.

### Statistics and reproducibility

Statistical analyses were carried out using Prism 7.0a (GraphPad). *P* values were calculated using the unpaired t-test or analysis of variance. All values are given as mean ± SEM. For animal studies, no statistical method was used to predetermine sample size. The experiments were not randomized.

## Acknowledgements

We are grateful to Ana Maria Cuervo (Albert Einstein College) for sharing KECs and Sirio Dupont (University of Padova) for sharing FLAG-TAZ WT and 4SA plasmids. We thank Nicolas Goudin from Image analysis center (SFR Necker INSERM US24, CNRS UMS 3633) for designing the macro used to analyze the Zebrafish immunofluorescence data. We thank Enrico Moro (University of Padova, Italy) for providing Tg(hsp70l:RFP-Rno.Map1lc3b) embryos, as well as Marion Delous and Sophie Saunier (Imagine institute) for sharing their Tg(wt1b:GFP) zebrafish line. Nicolas Kuperwasser is acknowledged for editing and proofreading. This work was funded by Institut National de la Santé et de la Recherche Médicale (INSERM), Centre National de la Recherche Scientifique (CNRS), Université Paris Cité and Agence Nationale de la Recherche (ANR; R18004KK, R16167KK and R18158KK to P.C.; R18176KK to N.D.).

## Author contributions

Conceptualization: A.C.-T., N.D. and P.C. Formal analysis: A.C.-T., F.R., M.G.-T. and A.R. Investigation and validation: A.C.-T., F.R., M.G.-T., A.R. and M.B. Resources: N.D., P.C., E.M. and F.T. Data curation: A.C.-T. and N.D. Writing (original draft): A.C-T, N.D. and P.C.; Writing (comments): E.M. and F.T. Visualization: A.C.-T., F.R. and A.R. Supervision: N.D. and P.C. Project administration and funding acquisition: N.D., P.C. and E.M.

