## Supplementary Table 1 for "YAP-dependent autophagy is controlled by AMPK, SIRT1 and flow intensity in kidney epithelial cells"

| siRNA name | Qiagen Cat # | Target sequence |
|---|---|---|
| Mm_Yap1_6 | SI02689113 | ACCCTTGAACATATACATTTA |
| Mm_Yap1_7 | SI02711205 | AACATCCTATTTAAATCTTAA |
| Mm_Wwtr1_6 | SI02697149 | CTGCATTTCTGTGGCAGATAA |
| Mm_Wwtr1_7 | SI02720599 | TTCCTTAATCACATAGAGAAA |
| Mm_Kif3a_2 | SI00175987 | ACGAACCTCCAAAGACATTTA |
| Mm_Kif3a_4 | SI00176001 | GACCCAGAGGTTAGAGGTTAA |
