## Supplementary Table 2 for "YAP-dependent autophagy is controlled by AMPK, SIRT1 and flow intensity in kidney epithelial cells"

| Plasmid | Source | Identifier |
| --- | --- | --- |
| pCMV-FLAG-YAP-5SA | Addgene | #27371 |
| pCMV-FLAG-YAP-5SA/S94A | Addgene | #33103 |
| pDONR221-P5P2-FLAG-YAP1 | Addgene | #79503 |
| pDONR221-P5P2-FLAG-YAP1-S61A | This study | |
| pCMV-mRFP-FLAG-YAP1 | This study | |
| pCMV-mRFP-FLAG-YAP1-S61A | This study | |
| pFLAG-mTAZ | Dupont S. | PMID: 21654799 |
| pFLAG-mTAZ-4SA | Dupont S. | PMID: 21654799 |
| 8xGTIIC-luciferase | Addgene | #34615 |
| pRL-TK-Renilla-Luciferase | Promega | #E2241 |
