## Supplementary Table 3 for "YAP-dependent autophagy is controlled by AMPK, SIRT1 and flow intensity in kidney epithelial cells"

| Target | Forward | Reverse |
|---|---|---|
| *Ankrd1* | CTGTGAGGCTGAACCGCTAT | CCAGTGCAACACCAGATCCA |
| *β-actin* | GGCCAACCGTGAAAAGATGA | ACCAGAGGCATACAGGGACAG |
| *Bcl2* | GTGGATGACTGAGTACCT | CCAGGAGAAATCAAACAGAG |
| *Cptp* | GTGGAAGGAAACTAGGCCCC | CCAGTGGAAAAGCGTAGGGT |
| *Cyr61* | AGAGGCTTCCTGTCTTTGGC | CCAAGACGTGGTCTGAACGA |
| *Ptplad2* | GTCGAACAGAGCAGGAGGAAAC | CATTTGCTGTTGCCCAAGGAAT |
| *Ppargc1a* | TAAACTGAGCTACCCTTGG | CTCGACACGGAGAGTTAAAGGAA |
| *Rubicon* | CTCTGAGCAAGACTTTGGCAGC | GCACTTCATCAGCTCAATGGCG |
| *Tfam* | CCGAAGTGTTTTTCCAGCAT | GCGTGCAATTTTCCTAACCA |
